# Manipulating single-unit theta phase-locking with PhaSER: An open-source tool for real-time phase estimation and manipulation

**DOI:** 10.1101/2023.02.21.529420

**Authors:** Zoé Christenson Wick, Paul A Philipsberg, Sophia I Lamsifer, Cassidy Kohler, Elizabeth Katanov, Yu Feng, Corin Humphrey, Tristan Shuman

**Author notes:** co-first authors (Zoé Christenson Wick, Paul A Philipsberg).

## Abstract

The precise timing of neuronal spiking relative to the brain’s endogenous oscillations (i.e., phase-locking or spike-phase coupling) has long been hypothesized to coordinate cognitive processes and maintain excitatory-inhibitory homeostasis. Indeed, disruptions in theta phase-locking have been described in models of neurological diseases with associated cognitive deficits and seizures, such as Alzheimer’s disease, temporal lobe epilepsy, and autism spectrum disorders. However, due to technical limitations, determining if phase-locking causally contributes to these disease phenotypes has not been possible until recently. To fill this gap and allow for the flexible manipulation of single-unit phase-locking to on-going endogenous oscillations, we developed PhaSER, an open-source tool that allows for phase-specific manipulations. PhaSER can deliver optogenetic stimulation at defined phases of theta in order to shift the preferred firing phase of neurons relative to theta in real-time. Here, we describe and validate this tool in a subpopulation of inhibitory neurons that express somatostatin (SOM) in the CA1 and dentate gyrus (DG) regions of the dorsal hippocampus. We show that PhaSER is able to accurately deliver a photo-manipulation that activates opsin+ SOM neurons at specified phases of theta in real-time in awake, behaving mice. Further, we show that this manipulation is sufficient to alter the preferred firing phase of opsin+ SOM neurons without altering the referenced theta power or phase. All software and hardware requirements to implement real-time phase manipulations during behavior are available online (https://github.com/ShumanLab/PhaSER).

## INTRODUCTION

Neural oscillations, and particularly hippocampal theta, have long been theorized to organize neural activity – chunking sensorimotor information into discrete bins that facilitate the processing and storage of that information (Bures and Fenton, 2000; Buzsáki, 2002; Colgin, 2011). Support for this notion lies in the fact that the vast majority of well-defined cell subpopulations seem to lock their spiking to a certain phase (i.e., the rising phase, peak, falling phase, or trough) of each theta cycle. For example, hippocampal inhibitory neurons, which are highly heterogeneous, show subtype-specific oscillation-locked activity (Klausberger et al., 2003; Klausberger and Somogyi, 2008; Somogyi et al., 2013; Dudok et al., 2021). Axo-axonic cells, for instance, increase their firing selectively at the peak of theta (when referenced to the CA1 pyramidal layer LFP) while dendritically targeting somatostatin-expressing (SOM)+ cells increase their firing near the trough of theta. In other words, these two inhibitory cell populations release GABA onto their postsynaptic targets at opposing times of the theta cycle (near the peak versus near the trough) and also onto different anatomical sub-compartments (the axon initial segment versus the dendritic arbor). It is therefore thought that they act to gate the computational processing of their postsynaptic targets in distinct ways during these two theta phases (Somogyi et al., 2013). This idea is strongly supported by evidence from correlational and computational studies that suggest that inhibitory phase-locking is a powerful regulator of hippocampal function (Cutsuridis and Hasselmo, 2012; Cutsuridis and Poirazi, 2015) and that information encoding and retrieval may occur at opposing phases of the hippocampal theta cycle (Hyman et al., 2003; Kwag and Paulsen, 2009; Siegle and Wilson, 2014).

Indeed, the extent to which single-units organize their spiking within the theta oscillation (i.e., the strength of their theta phase-locking) during learning is correlated with one’s likelihood to correctly recall that information later (Rutishauser et al., 2010). Further, rodent models of diseases with cognitive dysfunction show disrupted theta phase-locking (Lenck-Santini and Holmes, 2008; Sigurdsson et al., 2010; Bender et al., 2016; Lopez-Pigozzi et al., 2016; Lazaro et al., 2019; Shuman et al., 2020). Computational models emphasize the importance of inhibition in information processing in the hippocampus, suggesting that the successful theta phase-locking of DG inhibitory neurons may directly gate information flow through the DG (Cutsuridis and Poirazi, 2015) as well as influence network connectivity in CA1 (Cutsuridis and Hasselmo, 2012). Together, this suggests that theta phase-locking of hippocampal inhibitory neurons may control information processing and gate hippocampal functions. Yet, despite much theoretical and correlative support for this hypothesis, there is a lack of direct causal evidence that inhibitory theta phase-locking is critical for the processing of information in the hippocampus.

This gap in knowledge is due, in part, to the highly technical and challenging nature of manipulating phase-locking. To experimentally manipulate phase-locking, one must detect the oscillatory phase in real-time in awake, behaving animals while perturbing neural activity in a phase- and cell type-specific manner. Fortunately, several recent advances have made substantial progress in enabling real-time closed-loop manipulations of network activity relative to theta phase. Indeed, optogenetic activation of parvalbumin (PV)+ neurons at the peak and trough of theta had distinct effects on behavior when the manipulation occurred during information encoding versus retrieval (Siegle and Wilson, 2014). Oscillation-driven closed-loop manipulations have also been utilized to determine the influence of gamma power on memory consolidation (Kanta et al., 2019), to test how phase-specific manipulations may induce plasticity of both network and behavioral responses (Desideri et al., 2018; Gordon et al., 2022), and to test whether phase-locked reactivation of memory-related principal cell ensembles impacts the behavioral expression of that memory (Rahsepar et al., 2022). Additionally, new open-source toolkits, such as TORTE and CLoSES, have been developed to flexibly deliver oscillation-locked manipulations (Zelmann et al., 2020; Schatza et al., 2022) and can be utilized for a wide variety of applications, from rodent LFP to human EEG recordings. Notably, however, while all of these studies utilize phase-specific manipulations, none have monitored the ability of these manipulations to alter the phase-preference of single-units during behavior.

Here, we describe and validate PhaSER (<u>Pha</u>se-locked <u>S</u>timulation to <u>E</u>ndogenous <u>R</u>hythms), an open-source, user-friendly platform that allows for fast, flexible, and highly accurate closed-loop delivery of perturbations at any specified phase of an oscillation. We find that PhaSER estimates theta phase with outstanding accuracy across a range of physiological theta powers in animals while they traverse a virtual linear track and we validate the utility of this tool for precisely manipulating the theta phase preference of a subtype of inhibitory neurons – somatostatin-expressing (SOM+) neurons in the dorsal hippocampus.

## RESULTS

### PhaSER performs precise and accurate theta phase-estimation in awake, behaving animals

To test PhaSER’s ability to alter the theta phase preference of SOM+ neurons in the hippocampus, we first expressed an excitatory opsin, channelrhodopsin (ChR2), in our neurons of interest. To do this, we injected a Cre-dependent ChR2 (AAV1-hSyn-DIO-hChR2(H134R)-eYFP) into the dorsal CA1 and DG of male and female SOM-Cre mice. At the same time, we implanted a stainless steel headbar on the skull to allow head-fixation. After recovery from surgery, we trained these mice to traverse down a virtual linear track to earn water rewards (**Fig 1A**; see <u>Methods</u> for details).

**FIGURE 1.**
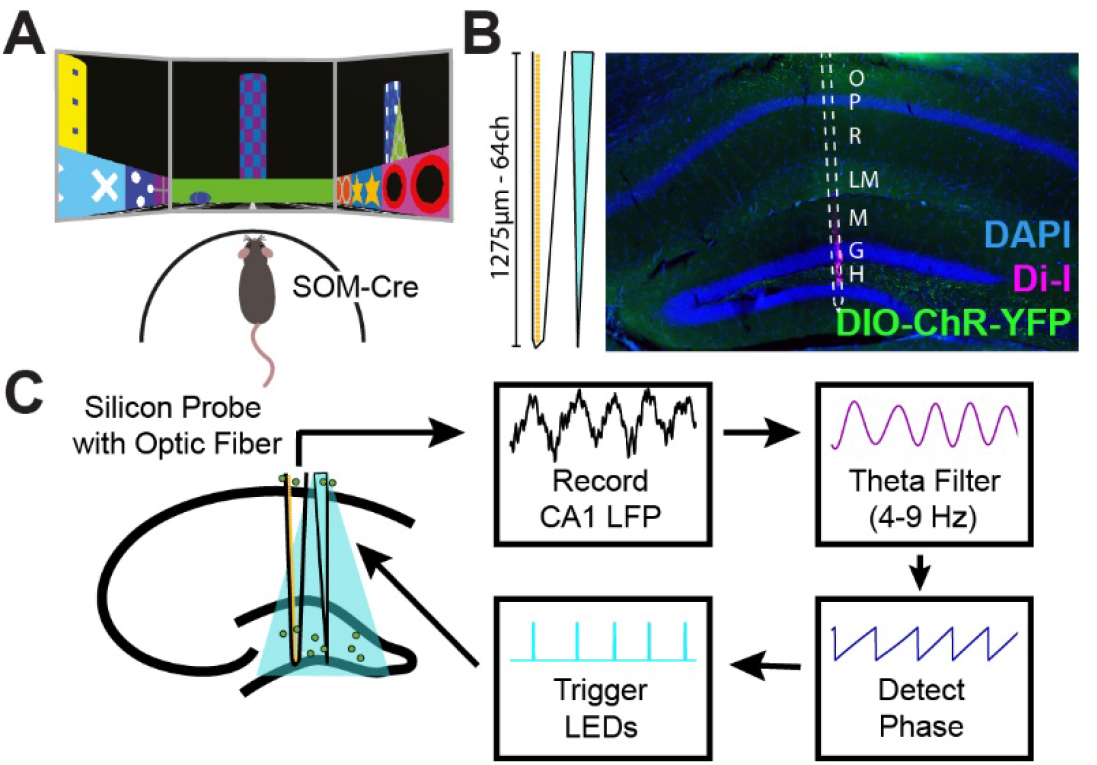
Acute silicon probe recordings for real-time theta phase detection and phase-locked optogenetic manipulations using PhaSER. **A)** Somatostatin (SOM)-Cre mice expressing Cre-dependent channelrhodopsin (ChR) are trained to navigate a virtual linear track while headfixed on a spherical treadmill. **B)** 64-channel acute silicon probe with tapered optic fiber is coated with dye (DiI, magenta) and lowered into the dorsal hippocampus, with electrodes spanning all layers of CA1 and the DG. Cre-dependent channelrhodopsin is injected into the dorsal CA1 and DG, expressing in SOM+ inhibitory neurons in both regions. **C)** Simplified schematic of PhaSER’s signal processing pipeline. A theta reference channel is selected from the pyramidal cell layer of CA1 and the raw local field potential (LFP) is passed through a real-time theta filter before using a Hilbert transform for theta phase detection. When the detected theta phase matches the target phase, a blue LED is triggered to turn on and activate the ChR expressed in SOM+ neurons of the dorsal hippocampus.

Once fully trained, we performed an acute silicon probe recording where a dye-coated 64 channel artifact-free silicon probe (H3, Cambridge Neurotech) with an attached tapered optic fiber was slowly lowered into the dorsal hippocampus, with electrodes spanning the CA1 and DG regions (**Fig 1B**). We first collected a baseline period of at least 50 laps on the virtual linear track before any photo-manipulations were applied. At the end of this baseline, we photo-tagged SOM+ neurons using a 1Hz 10ms blue light stimulation paradigm. This photo-tagging period allowed us to follow the activity of opsin+ neurons across both the baseline and phase-specific manipulation periods.

To test if we could accurately deliver phase-specific manipulations (i.e., peak-targeted or trough-targeted stimulations; see **Fig 1C** for a simplified schematic), we referenced theta from an electrode in the deep pyramidal layer of CA1 and trained the phase-estimation algorithm on approximately 2 minutes of baseline data. Phase estimation was performed as follows: the raw data from the pyramidal cell layer reference electrode was passed into a NI-DAQ where it was zero-phase filtered for theta (4-9Hz; **Fig 2**). Filtering in real-time introduces distortions at the edges of the sampling window which can negatively impact the accuracy of real-time phase estimation, so the edges were trimmed and an established autoregression model (Chen et al., 2013) was used to predict approximately one theta cycle into the future (**Fig 2A**). A Hilbert transform was then applied on the predicted theta oscillation and a 5ms blue light pulse was triggered when the target phase was detected. To limit our phase-locked manipulation to periods of relatively strong theta power, we only delivered this manipulation when the mouse was running on the virtual track (Wyble et al., 2004). Because we aimed to manipulate theta phase-locking of single-units in real-time, it was important to minimize the lag from data acquisition to signal processing to light trigger. With our hardware setup (**Fig 2B**, see <u>Methods</u> for details), the signal processing latency is ∼3ms, which can be offset by targeting a phase ∼9º prior to the desired stimulation phase.

**FIGURE 2.**
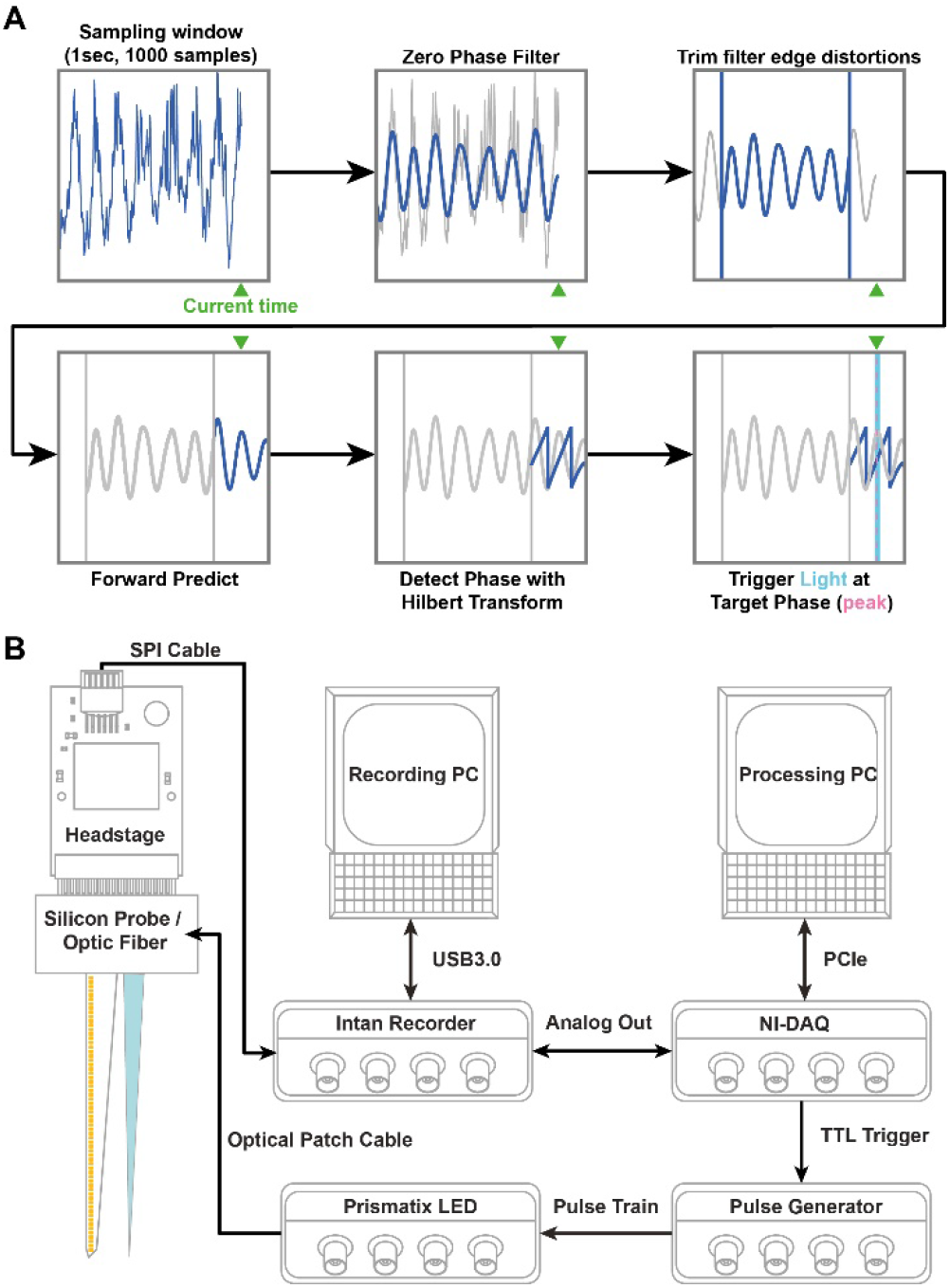
High-speed hardware and an autoregressive model allow PhaSER to predict and detect the theta phase in near real-time. **A)** A 1 second window of data is down-sampled to 1000Hz and is then zero-phase filtered and the first and last 150ms of the filter window are trimmed to remove edge distortions. An autoregressive forward prediction model is then used to extrapolate the trimmed 150ms followed by an additional 150ms forward prediction. Finally, the Hilbert transform is applied and the current phase is taken at the center of the 300ms forward prediction window (i.e., the current time). When the estimated current phase matches the target phase (e.g., the theta peak), light delivery is triggered. **B)** Electrophysiological data is digitized and amplified at the headstage before being passed into an Intan Recording System and saved to a Recording PC. The theta reference channel is passed from the Intan Recorder as an Analog signal to a National Instruments DAQ (NI-DAQ). The signal is transferred to a processing PC via high-speed PCIe connector before undergoing phase estimation as described in **A**. Upon detection of the target phase, the NI-DAQ sends a TTL trigger to a pulse generator, which switches on a LED connected via patch cable to the implanted tapered optic fiber. To achieve low-latency (∼3ms) signal processing, we use high-speed SPI and PCIe cables to swiftly carry the electrophysiological signal from the silicon probe to the processing PC. Additionally, to prevent overburdening the CPU, we use separate PCs for recording and real-time signal processing, though this may not be strictly necessary.

To ensure that PhaSER was performing accurate and precise real-time phase estimation, we compared the light delivery times during peak- and trough-stimulations to the offline filtered theta oscillation (**Fig 3**). To measure the precision and accuracy of phase-specific stimulation we calculated the magnitude of phase preference (r-value or mean resultant vector length) and the mean phase of stimulation (mu-value), respectively. We found that PhaSER was both highly precise (r=0.64 ± 0.04 for trough and r=0.66 ± 0.03 for peak, n=13 animals) and accurate (phase of trough triggers=188.3° ± 17.1° (circular mean ± standard deviation); peak triggers=11.5° ± 22.0°; **Fig 3C**). Additionally, to confirm that PhaSER was capable of performing precise phase-estimation across a range of physiological theta powers, we compared the theta power in the referenced channel during the baseline period to the trigger precision (r-value) of the light delivery during trough- and peak-targeted stimulation and found no significant correlation between the two variables (trough trigger precision vs baseline theta power: Pearson r = -0.03, *p*=0.93; peak trigger precision vs baseline theta power: r = 0.38, *p*=0.22, n=12 animals; **Fig 3D**).

**FIGURE 3.**
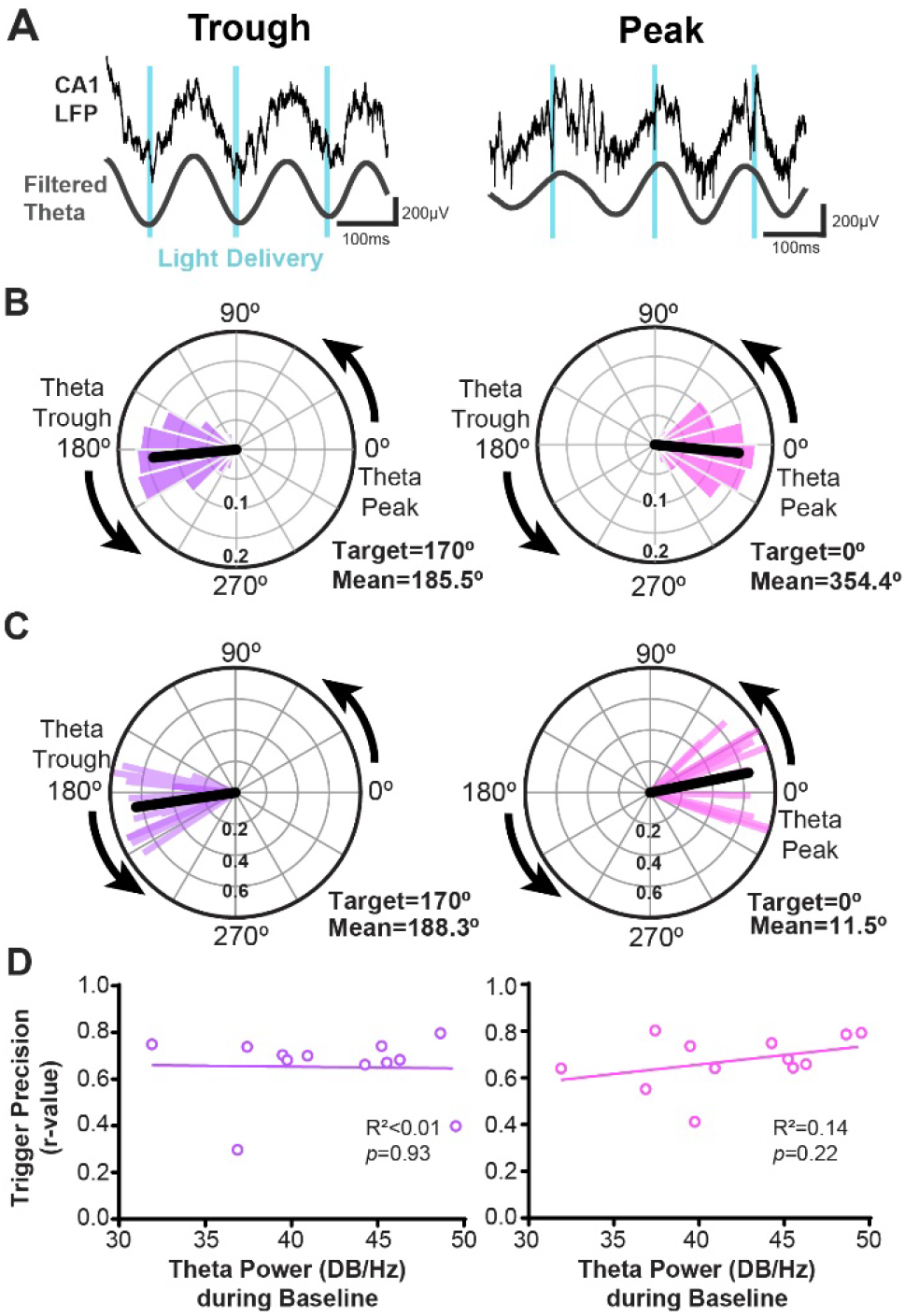
PhaSER accurately detects and triggers light delivery to a specified theta phase. **A)** Light delivery (blue) occurs at the trough (left) or peak (right) of CA1 theta (both the raw reference LFP and offline filtered signal are shown). **B)** Circular distribution of light pulses in a representative animal targeted to the theta trough (left, target 170°) are centered around the theta trough (mean phase of light delivery = 185.5°) and light pulses targeted to the theta peak (right, 0°) are centered around the theta peak (mean phase of light delivery = 354.4°, or -5.6°). The mean phase is shown in black. **C)** Polar plot showing the distribution of the mean phase of light delivery across mice when targeting the trough (left, target 170°, mean = 188.3°) and peak (right, target 0°, mean = 11.5°) of CA1 theta. The trigger precision (r-value, length of the resultant vector) is shown on the radial axis. **D)** The trigger precision (r-value) targeted towards the trough (left) or peak (right) is not significantly correlated with theta power during baseline (trough: Pearson r=-0.03, p=0.93; peak: Pearson r=0.38, *p*=0.22, n=12 animals).

### Real-time phase-specific manipulations shift the theta-phase preference of SOM+ neurons in awake, behaving animals

Once we confirmed that PhaSER was delivering light accurately to the targeted theta phases, we examined whether this phase-specific manipulation was able to shift the theta phase preference of opsin+ neurons. Single-units were identified and clustered with Kilosort2.5 (Stringer et al., 2019; Steinmetz et al., 2021) and inspected and curated using Phy2 (Steinmetz et al., 2021). The response of these clustered units to 100 pulses of blue light (10ms duration, 1Hz) was then assessed. Clusters showing a significant (α<0.001) increase in firing within 4ms of the light onset were considered photo-tagged single-units and are referred to as SOM+ neurons from here on (**Fig 4A-B**). Once identified, we examined the baseline phase preference of these SOM+ neurons and found they had a moderate preference (r-values 0.25 ± 0.05, n=19 cells) for firing near the trough of the CA1 theta oscillation (**Fig 4C-D**; mu-values 152.4° ± 42.5°, n=19 cells).

**FIGURE 4.**
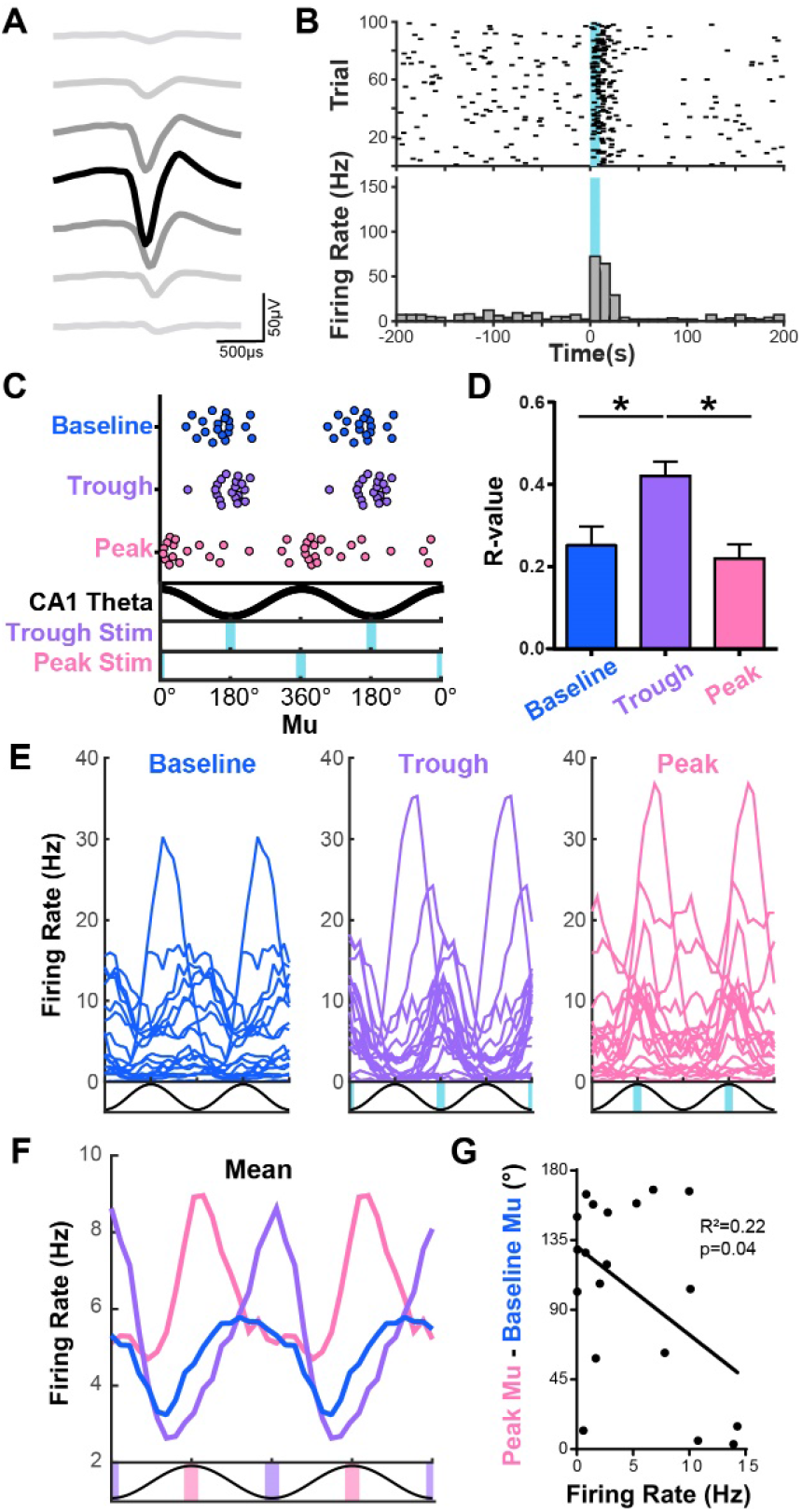
Phase-specific manipulations alter the phase-preference of opsin-expressing neurons. **A)** Average waveform of a photo-tagged neuron with waveform shown across 7 channels, centered around the channel with the highest spike amplitude. **B)** Raster plot and histogram of a photo-tagged neuron showing a significant increase in spiking aligned to light onset (10ms light pulse). **C)** Preferred firing phase (mu) of photo-tagged putative SOM+ neurons during baseline (blue), trough-targeted stimulation (purple), and peak-targeted stimulation (pink). Peak-targeted stimulation significantly shifts the distribution of preferred firing phase compared to baseline or trough-targeted stimulation periods (Kuiper test *p*<0.05). **D)** The strength of phase preference, or r-value, was significantly increased during trough-targeted stimulation compared to baseline or peak-targeted stimulation periods (repeated measures ANOVA with *posthoc* Holm-Sidak tests, *p*<0.05). **E-F)** Firing profiles relative to CA1 theta of individual photo-tagged SOM+ neurons during baseline, trough-, and peak-targeted stimulation periods with mean firing profiles shown in **F. G)** The difference in opsin+ SOM neurons’ phase preference (mu) during peak stimulation compared to their baseline preference is negatively correlated with the cells’ baseline firing rate (Pearson r = -0.47, *p*=0.04).

To test if trough- or peak-targeted stimulation altered the phase preference of the opsin+ cells, we then compared their mean preferred firing phase (mu-value) during baseline to their mean preferred firing phase during these manipulation periods. We found that trough-targeted stimulation subtly shifted the mean preferred firing phase of SOM+ neurons closer to the theta trough (mu 180.0° ± 33.8°), but this comparison only approached significance (Watson-Williams test comparing baseline mu to trough mu, F=4.07, *p*=0.051, Kuiper test *p*>0.1, n=19 cells from 5 animals; **Fig 4C**). Additionally, trough-targeted stimulation significantly enhanced the magnitude (r-value) of opsin+ SOM neurons’ theta phase preference (**Fig 4D**; r-value 0.42 ± 0.04; repeated measures ANOVA showed a significant effect of Manipulation, *p*<0.0001, F=15.33, n=19 cells; with *posthoc* Holm-Sidak’s multiple comparisons tests showing a significant increase in r during trough-targeted stimulation compared to either baseline or peak-targeted stimulation, *p*<0.05).

Furthermore, peak-targeted stimulation was able to significantly shift the phase preference of these opsin+ SOM neurons toward the peak of CA1 theta (**Fig 4C**; mu 32.2° ± 57.0°; Watson-Williams test comparing baseline mu to peak mu, F= 34.03, *p*<0.0001; Kuiper test baseline vs peak: *p*=0.002, trough vs peak: *p*=0.001; n=19 cells) while maintaining a similar magnitude of phase-locking (**Fig 4D**; r-value 0.22 ± 0.03; Holm-Sidak *posthoc* test showing no significant difference between baseline and peak-stimulation, *p*>0.05, n=19 cells).

### Baseline firing frequency is negatively correlated with the malleability of theta phase-preference

While the majority of opsin+ SOM neurons’ mu values shifted toward the theta peak during peak-targeted stimulation, a few cells did not show fully shifted phase preferences (**Fig 4C**). Upon investigation of the individual firing profiles of the SOM+ neurons during baseline, trough-, and peak-targeted stimulations (**Fig 4E-F**), we noticed that the cells with higher average firing rates during baseline seemed less modulated by the peak-targeted manipulation. To test this, we compared the mean firing rate of each cell during baseline to the difference between that cell’s mu value during peak stimulation versus baseline. We found that the neurons’ firing rates were negatively correlated with how malleable their mu values were during peak-targeted stimulation (Pearson r=-0.47, *p*=0.04, n=19 cells, **Fig 4G**), with faster firing cells maintaining their initial phase preference while neurons with lower firing rates showed a greater change in mu between peak-stimulation and baseline periods. This may be due to the fact that while we were delivering stimulations to the peak of theta, the inputs driving these cells to fire near the trough remained intact. Thus, cells that are kinetically and metabolically capable of firing at faster frequencies may maintain their firing at their initially preferred phase in addition to being activated at the stimulation phase.

### Theta power and phase are not altered by phase-specific SOM+ cell manipulations

Some inhibitory neurons in the dorsal hippocampus have been shown to contribute to the local generation of theta, with stimulation of certain inhibitory cell populations changing the power and phase of the theta oscillation itself (Amilhon et al., 2015; Bezaire et al., 2016; Christenson Wick et al., 2019). Because altering the referenced theta oscillation with real-time manipulations may impact PhaSER’s ability to perform accurate phase-estimation, we tested whether applying trough- or peak-targeted stimulations to SOM+ neurons changed the power or phase of the referenced LFP. To look into this, we first tested for changes in power spectral density across all layers of dorsal CA1 and DG during trough- or peak-targeted stimulation compared to the baseline period (**Fig 5A**). We found no significant changes in power spectral density (multiple t-tests comparing power spectral density of oscillatory frequencies during trough- and peak-targeted stimulation periods to baseline [1-100Hz, 1Hz bins] across all 64 channels aligned by hippocampal layer, uncorrected *p*>0.05). This was further confirmed when we tested for changes in power specifically in the CA1 theta reference layer during baseline, trough-, and peak-targeted stimulations (**Fig 5B**; repeated measures ANOVA with Greenhouse-Gessier correction, F=1.82, *p*=0.25, n=5 animals). Finally, we tested if our stimulation reset the theta phase of the referenced channel, as such an effect would greatly diminish PhaSER’s ability to perform accurate phase-estimation (Wodeyar et al., 2021). To test for a theta phase reset with SOM+ neuron stimulation, we aligned the unfiltered, down-sampled reference LFP to 10ms light pulses (delivered at 1Hz frequency, see photo-tagging protocol) and averaged the LFP across 100 light pulses. We then compared the envelope of the averaged signal 250-300ms before to 250-300ms after the light delivery. If the stimulation of SOM+ neurons was resetting the theta phase, we would expect to see an increase in the mean LFP envelope lasting more than 1 theta cycle (i.e., >120ms). Instead, we saw no significant change in the mean envelope before versus after the aligned light pulses (paired t-test, *p*=0.27; **Fig 5C-D**). Together, this suggests that stimulation of SOM+ neurons is not significantly altering the theta power nor is it resetting the referenced theta phase.

**FIGURE 5.**
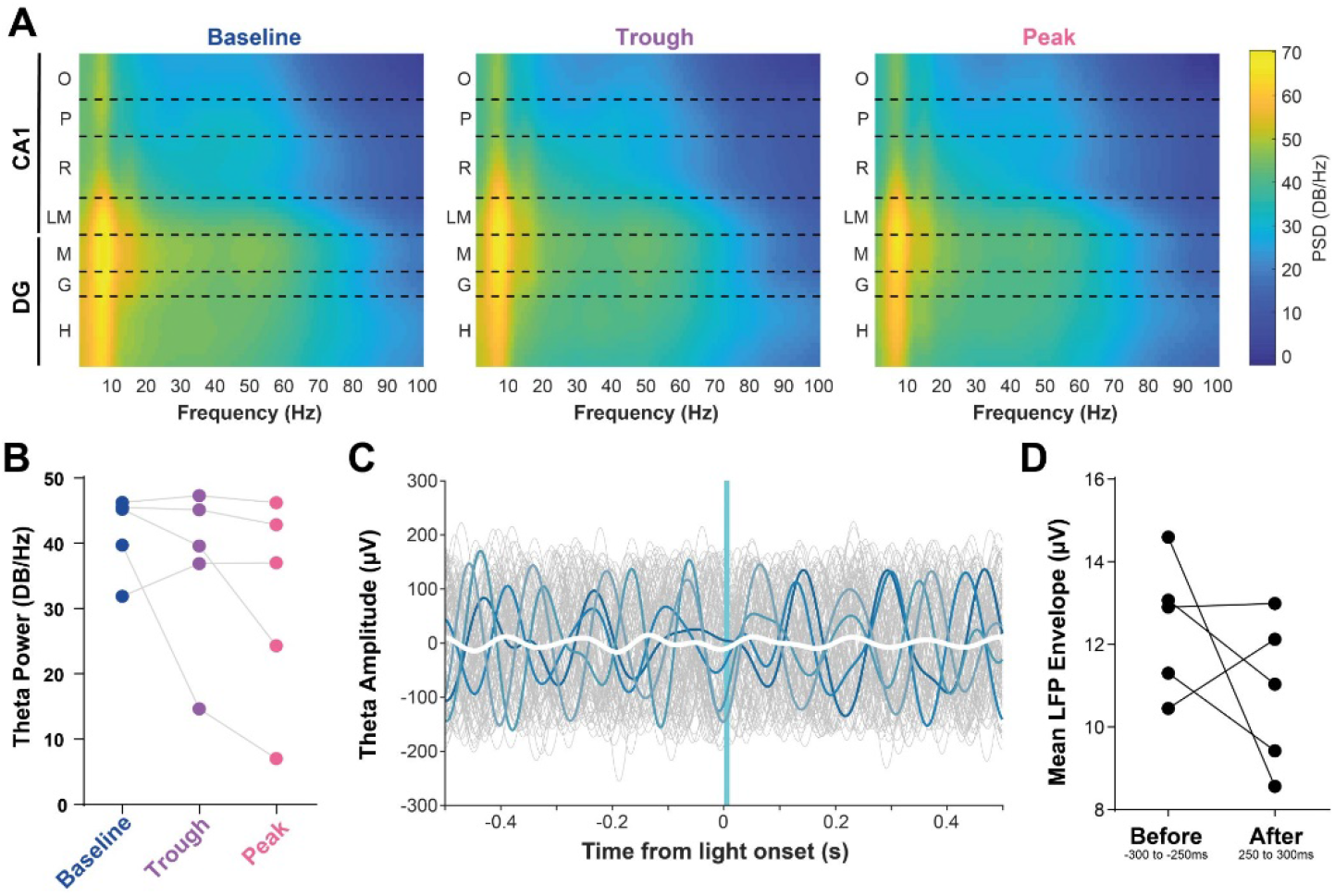
Theta phase-specific manipulation of opsin-expressing SOM+ neurons does not alter theta power or reset theta phase. **A)** Power spectral density (PSD) during running bouts of at least 3 seconds during baseline, trough-, and peak-targeted stimulation periods. Power while running during trough- and peak-stimulation periods was normalized to baseline for each animal and the distribution of changes in power was compared against zero in each channel across all frequencies (1Hz bins, 1-100Hz). No significant differences in PSD were found during trough- or peak-stimulation (uncorrected alpha of 0.05, n=5 animals). **B)** Theta power in the CA1 pyramidal reference layer was not significantly altered during trough- or peak-targeted stimulation (repeated measures ANOVA with Greenhouse-Gessier correction F=1.82, *p*=0.25, n=5 animals). **C)** Filtered theta from the CA1 pyramidal cell reference layer aligned to 100 light pulses (10ms duration). Randomly selected individual traces are colored in shades of blue, mean of all traces is in white. Note that this signal has been filtered for theta (5-12Hz) for visualization purposes, but analysis in **D** is on unfiltered data. **D)** The LFP envelope was calculated from down-sampled, unfiltered data aligned to light pulses as in **C** on a 50ms window starting 250ms before or 250ms after light delivery. Mean LFP envelope was compared using a paired t-test, *p*=0.27.

## DISCUSSION

Here, we introduce PhaSER, a new open-source tool for real-time phase-estimation and delivery of phase-specific manipulations. We show that it is capable of accurately and precisely targeting manipulations to specific phases of ongoing theta oscillations and that it can be used to manipulate the phase preference of SOM+ neurons in the dorsal hippocampus. This suggests that phase-specific manipulations can be a powerful tool to investigate the functional significance of theta phase-locking, a phenomenon that has been extremely challenging to causally interrogate in the past.

PhaSER can be flexibly utilized across a variety of recording platforms. With our hardware setup, the signal processing latency is approximately 3ms (or 9° in a 120ms theta cycle), which is more than sufficient to accurately deliver theta phase-specific stimulations and would, in theory, be fast enough to accurately target specific phases of gamma oscillations as well. Of course, the signal delay will differ based on the hardware used and therefore should be measured and adjusted for accordingly when applied in other setups. In cases where the signal delay negatively impacts the accuracy of manipulation delivery, the manipulation target phase can simply be set earlier in the oscillation cycle by an amount equivalent to the signal delay latency.

In addition to considering the hardware and signal latency when applying PhaSER to future studies, it will also be important to consider the spike kinetics, intrinsic excitability, and heterogeneity of any cell populations being manipulated in a phase-specific manner. For example, stimulating SOM+ neurons at the theta trough (near when they would typically fire anyways) did not significantly change the cells preferred firing phase (although this approached significance), but did increase the strength of their phase preference (i.e., r-value). In contrast, stimulations applied at the peak of theta, when these cells are normally silent, shifted the preferred firing phase but did not alter the strength of their phase preference compared to baseline. Furthermore, it seems that the ability of phase-specific stimulations to effectively shift the phase preference of SOM+ neurons was negatively correlated with each neuron’s baseline firing frequency. Therefore, this approach should be validated in each target cell population and bidirectional manipulations should be considered in cell populations with fast-spiking kinetics and strong excitatory inputs that would enable them to fire multiple action potentials within one theta cycle. To facilitate this, we have built multi-phase stimulation options into PhaSER such that an excitatory photo-manipulation could be delivered at one phase while an inhibitory photo-manipulation could be delivered at another phase. Importantly, while we have shown peak- and trough-targeted manipulations here, PhaSER is able to accurately target any specified phase of theta as its phase detection algorithm does not depend on identifying cycle maxima or minima.

We have shown that PhaSER’s phase estimation algorithm remains precise across a wide range of physiological theta powers; however, it is important to note that the phase estimation precision and accuracy will plummet when the referenced oscillatory power is greatly diminished, such as during the quiescent period after a seizure. The zero-phase filter will continue to find theta cycles in noisy or aperiodic activity (Donoghue et al., 2020; Wodeyar et al., 2021), therefore it is important to consider under what circumstances a manipulation should be triggered. Here, we have limited light deliveries to periods when the animal is running down a virtual linear track, thereby ensuring theta is reliable and highly prevalent during our manipulations (Wyble et al., 2004), which improves the accuracy and precision of our light deliveries. This will be of greater importance when applying phase-specific manipulations in the context of disease to determine if oscillation-driven manipulations can improve symptoms and drive long-term disease-modifying network reorganization. Phase-specific manipulations may be beneficial for the treatment of several diseases that are accompanied by rhythmopathies such as alterations in phase-locking, LFP power, cycle-to-cycle variability, oscillation shape, and cross-frequency coupling (Laurent et al., 2015; Shuman et al., 2017, 2020; Lazaro et al., 2019). In such situations, it will be especially important to validate the accuracy and precision of the phase-specific manipulations and consider how and when the manipulations may be more effectively applied.

Recent advances in electrical stimulation have led to new therapeutic options across a range of neurological and psychiatric disorders including epilepsy, depression, Parkinson’s disease, and obsessive-compulsive disorder (for reviews see Skarpaas and Morrell, 2009; Holtzheimer and Mayberg, 2011). Yet these stimulation patterns are often decoupled from endogenous brain rhythms and little is known about how the effectiveness of these stimulations may be improved by timing them to the brain’s ongoing activity. From both a clinical and basic-research perspective there is strong interest in acquiring a better understanding of how the timing of manipulations relative to endogenous oscillations and brain states impacts their effectiveness for treating psychiatric and neurological symptoms. By enhancing our knowledge of the precise impacts of phase-specific manipulations, we can inform how clinical stimulation strategies can complement, rather than compete with, the brain’s ongoing activity and improve these therapeutic interventions (Földi et al., 2021; Krause et al., 2022).

## METHODS

All experimental protocols were approved by the Icahn School of Medicine’s Institutional Animal Care and Use Committee, in accordance with the US National Institutes of Health guidelines.

### Animals

For experiments verifying the trigger accuracy, 7 male and 5 female SOM-Cre and PV-Cre mice were used. Both PV-Cre (Jackson Laboratory strain #017320; B6.129P2-Pvalb^tm1(cre)Arbr^/J; (Hippenmeyer et al., 2005) and SOM-Cre mice (Jackson Laboratory strain #013044; Sst^tm2.1(cre)Zjh^/J) were maintained homozygous with in-house breeding (Taniguchi et al., 2011). For optogenetic manipulation, 3 male and 3 female SOM-Cre mice were used. One animal was excluded from LFP analyses (**Fig 5**) due to having extremely low levels of opsin expression. Animals were housed in standard housing conditions (12-hour light, 12-hour dark) in the animal facility at the Icahn School of Medicine. Animals were group-housed with their littermates when possible or with an ovariectomized female (Jackson Laboratory strain# 000691; 129X1/SvJ) when littermates were not available or compatible. Animals were allowed *ad libitum* access to food and water except when water-restricted for training on the virtual linear track.

### Stereotaxic Surgery

The intrahippocampal virus delivery and headbar implant were performed during the same surgery under 1-3% isoflurane anesthesia in mice at 14 ± 1 weeks of age (97 ± 7 days). First, the skull was exposed and a small burr hole was drilled above the viral delivery site: 2mm posterior to bregma and 1.35mm to the right of bregma. A Nanoject III (Drummond Scientific Company) with glass capillary loaded with a Cre-dependent channelrhodopsin virus (ChR2; AAV1-EF1a-double floxed-hChR2(H134R)-EYFP-WPRE-HGHpA; Addgene cat #20298; 1.2×10^13^ GC/mL) was lowered into the burr hole until the tip was 2.1mm ventral to bregma. There, 120nL of virus was injected into the dentate gyrus at a rate of 2nL/s. Once the full 120nL was injected, the syringe was left for 3 minutes before being raised to CA1 (1.5mm ventral from bregma). There, an additional 120nL of virus was injected in the same manner.

After viral infusion, a stainless steel headbar was fixed onto the skull of the mouse. Lidocaine (2%, ∼0.03mL) was first injected subcutaneously over the skull, and the epaxial muscles along the neck were cleared from the skull’s surface. The mouse skull was thoroughly scored and then was stereotactically aligned to the headbar before the headbar was fixed near the surface of the skull with cyanoacrylate glue and dental cement (Lang Dental). Dental cement was built up to create a well around the exposed skull which was then filled with Kwik-Sil (World Precision Instruments). Once dried, the Kwik-Sil was then covered with a final layer of dental cement. Meloxicam (5mg/kg) or Carprofen (5mg/kg) was administered subcutaneously during and for 2 days following surgery together with a 7-day course of ampicillin (20mg/kg). Animals were returned to the animal housing facility after recovering from anesthesia on a heating pad.

### Virtual Reality (VR) Training

In the days following surgery, while postoperative drugs were administered, mice were gently handled for 5 minutes daily. Once recovered from the surgery and fully habituated to being handled (∼3 days), mice were introduced to head-fixation while being allowed to explore a flat surface for 5 minutes daily until they demonstrated the ability to move while head-fixed (∼3 days). Mice were then water restricted and maintained at a body weight of around 82% of their initial weight while they continued the rest of their daily training (as in Shuman et al., 2020). During water restriction, mice were weighed and monitored daily for signs of dehydration, fatigue, or infection and were immediately given free access to water if any signs of distress were observed. After water restriction began, mice were introduced to head-fixation while standing/walking on a spherical treadmill locked to rotate on only one axis. Once the mice had gained coordination and strength on the spherical treadmill (usually ∼3 days of training for 10-20 minutes per day), mice were trained to lick from a water port that delivered a 4µL drop of water per lick. Once the mice had learned to reliably earn water from the water port, they were trained to run along virtual linear tracks of increasing length where water rewards would be delivered at the end of the track before they were teleported back to the beginning of the track. The virtual linear track, which was created with ViRMEn, an open-source MATLAB-based software package (Aronov and Tank, 2014), was displayed across three flat monitors angled around the front of the spherical treadmill. Once the mice were consistently earning their daily water in 1 hour or less on the longest (∼2 meter) virtual track while maintaining their body weight at ∼82%, they were prepared for acute silicon probe recording. Virtual track training took 5-8 days.

### Craniotomy & Ground Implantation Surgery

One day prior to an acute silicon probe recording, a craniotomy and ground implantation was performed. First, the most superficial layer of dental cement was drilled off and the Kwik-Sil was removed, exposing the skull. Then a burr hole was drilled above the left hemisphere of the cerebellum, and an Ag/AgCl-coated silver reference wire (Warner Instruments) was slipped between the skull and dura and fixed with cyanoacrylate glue and dental cement. A 1.5mm diameter craniotomy was also drilled over the right hippocampus at this time (centered on the following coordinates: 2mm posterior, 1.45mm right from bregma). The craniotomy was covered with buffered artificial cerebrospinal fluid (ACSF; in mM: 135 NaCl, 5 KCl, 5 HEPES, 2.4 CaCl_2_, 2.1 MgCl_2_, pH 7.4; (Cai et al., 2016; Shuman et al., 2020) and the exposed skull was again covered with Kwik-Sil. Mice were returned to their home cages overnight after recovering on a heating pad.

### Acute Silicon Probe Recordings & Recording Hardware

The day following craniotomy, mice were set up for acute silicon probe recordings (15.5 ± 0.5 weeks of age (109 ± 4 days of age; 19 ± 1 day after initial surgery)). First, a 64 channel Cambridge NeuroTech (ASSY-77-H3) silicon probe (single shank, 8mm length, sharpened tip) with attached lambda-b optic fiber 100µm core (tapered 1.2mm) was painted with dye (Invitrogen, Vybrant DiI, V22885) so the probe track could be visualized later. Mice were then head-fixed on the spherical treadmill and the Kwik-Sil covering the skull was replaced with a buffered ACSF solution. Head-flat skull positioning in the dorsal-ventral and medial-lateral directions was then established to within 50µm (bregma-to-lambda) before the silicon probe was lowered into the right dorsal hippocampus at the following approximate coordinates: 2mm posterior, 1.45mm lateral, 2.1mm ventral to bregma. After reaching the target depth and confirming probe location with electrophysiological signatures of the dorsal hippocampus, mineral oil was placed over the buffered ACSF solution and the probe was allowed to settle for 1 hour before starting the virtual linear track and recording (Shobe et al., 2015; Shuman et al., 2020).

Electrophysiological signals were sent to an Intan headstage (RHD 64-Channel Recording Headstage, Intan Technologies) via a Samtec to Omnetics adaptor (ADPT A64-Om32×2, Cambridge NeuroTech) for pre-amplification and digitization. An Intan recording controller (RHD2000 Intan 1024ch Recording Controller, Intan Technologies) collected and logged signals at a 25kHz sampling rate from each electrode channel as well as from analog inputs reflecting the speed of the spherical treadmill, position on the virtual linear track, time of water delivery, time of the mouse’s licking behavior, and blue light trigger and delivery times. Because the Intan recording controller logs data to the PC over a USB connection, which is relatively slow, we selected a single electrode channel in the CA1 pyramidal cell layer (as identified by its location relative to a stereotypical theta phase shift and higher density of spikes) to route as an analog output going to a separate, low-latency, PCIe data acquisition device (NI-DAQ PCIe-6321, National Instruments).

Once the recording began, mice ran down the 2m virtual linear track at least 50 times to collect a baseline before any photo-manipulations were introduced. Immediately following this baseline period, single-units were photo-tagged with 100 1Hz square pulses (10ms duration) of blue light (LED, 455nm, Prizmatix, 1mW) sent via a Pulser Plus (Prizmatix) pulse train generator. Five minutes following the photo-tagging procedure, a baseline of at least 120s was acquired to train the autoregressive model for phase estimation (described in detail below). Photo-stimulation locked near the trough (170°) or peak (0°) was then delivered for at least 25 trials during locomotion. No phase-specific photo-stimulations were delivered during rest. The order of trough- and peak-targeted stimulations was counterbalanced across animals with at least 5 minutes between the two manipulation periods.

### Real-time Signal Processing & Phase-Specific Manipulation

In order to apply blue light stimulation locked to the ongoing theta oscillation in real-time, we created a custom LabVIEW program, which can be found on our lab’s github (https://github.com/ShumanLab) along with user-friendly documentation. The 25kHz recorded signal from the CA1 pyramidal layer reference channel was first down-sampled to 1kHz. Then, in order to do theta phase-specific manipulations, we filtered the down-sampled signal for the theta band (4-9Hz) in real-time. Because digital filters introduce phase lags and distortions, a zero-phase filter (first-order Butterworth) was employed. Zero-phase filtering is a technique in which a digital filter (in this case, an infinite impulse response (IIR) filter), which introduces a phase lag, is applied first in the forward direction and then in the reverse direction to cancel out all phase shifts in the output signal. To extract the phase from this filtered signal, we used the Hilbert transform. The Hilbert transform can be used to calculate the phase angle of a real signal at every point in time, however both digital filters and the Hilbert transform introduce edge effects – distortions at the edge of the analysis window. To circumvent these distortions, and therefore avoid incorrectly identifying the theta phase, we removed 150 samples from the beginning and end of the filtered signal within the 1000 sample sliding filter window and then used a 13^th^ order auto-regressive model that has been described previously (Chen et al., 2013). The auto-regressive model was trained on ∼2 minutes of baseline data using the Yule-Walker method. This model was then used to reconstruct the 150 most recent datapoints without edge effects and then forward predict an additional 150 samples (which equals approximately 1 theta cycle). This placed the current time point at the center of the 300-sample prediction window. The Hilbert transform was then applied to the resulting signal and the current phase angle was calculated from the center of the phase-estimation window, which corresponds to the real-time signal without edge effects. A schematic of this process is shown in **Fig 2A**. To limit photo-stimulation to periods when the animal was locomoting, a movement threshold was set based on the analog signal from the spherical treadmill. When this value passed the threshold indicating locomotion, and the Hilbert transform phase angle crossed the threshold of the light delivery target phase, a TTL trigger was generated by the NI-DAQ and passed to a pulse generator (Pulser Plus, Prizmatix) which was programmed to deliver 10ms of blue light (LED, 455nm, Prizmatix, 1mW) at the target phase. The theta trough stimulation target was 170° and the theta peak stimulation target was 0°. The latency of this real-time signal processing system from electrode to light trigger is approximately 3ms.

### Histology

Following recording, the silicon probe was removed from the brain and the mice were deeply anesthetized with isoflurane (5%) before being decapitated. The brains were quickly dissected and dropped in 4% paraformaldehyde where they stayed for ∼24 hours. 50µm thick coronal sections were prepared in 1x phosphate buffered saline (PBS; Fisher BP399) using a vibratome (Leica VT1000S). All sections containing hippocampus were then mounted for fluorescent imaging with DAPI (SouthernBiotech DAPI Fluoromount-Gm, 0100-20). Sections were later imaged to confirm viral expression (visualized via eYFP) as well as to determine probe location (DiI). Images were taken using a Leica DM6B fluorescence microscope equipped with a Lumencor Light Engine and Leica DFC9000 GT camera. Hippocampal anatomical locations of the probe tracks and stereotaxic coordinates were compared to the electrophysiological signals and a mouse brain atlas (Franklin and Paxinos, 2007).

### Post-Processing & Analysis of Single-Unit and Local Field Potential Data

All data analysis was performed with custom scripts using MATLAB 2017a. Data files were first concatenated into continuous signals for each channel. To extract single-units, all channels were highpass filtered and background subtracted for clustering into putative single-units using Kilosort2.5 (Stringer et al., 2019; Steinmetz et al., 2021) and Phy2 (Steinmetz et al., 2021). This automated spike sorting pipeline reliably isolates single-units using a template-matching approach that is iteratively updated and includes drift correction, a critical step for stable cell isolation across the entire recording. Once single-units were isolated using Kilosort2.5 and manually confirmed in Phy2, we tested each single-unit for its response to our photo-tagging stimulation protocol. To identify photo-tagged cells, we built a spike probability distribution based on the 500ms prior to each light stimulation (which occurred at a frequency of 1Hz). If a single-unit’s spike frequency during the first 4ms of light delivery was significantly above the distribution mean (where α = 0.001), then the cell was classified as opsin+. Opsin+ cells were assessed for their phase preference during the baseline period as well as their response to the trough and peak phase-specific manipulations. Phase-locking was quantified by an r-value (magnitude of phase preference) and a mu-value (mean phase of firing) relative to the deepest pyramidal layer channel (i.e. the pyramidal cell layer channel closest to the stratum oriens).

For local field potential (LFP) analyses, signals were down-sampled to 1kHz, bandpass filtered (theta: 5-12Hz) using zero-phase digital filtering and LFP power was compared using the Chronux library (Mitra and Bokil, 2008; Mitra et al., 2018). Power spectral density (PSD) was calculated from the unfiltered, down-sampled LFP at each electrode position during running bouts that were at least 3 seconds in duration. A spectrogram was constructed by taking the mean PSD across running bouts within each frequency bin (1Hz bin size) and electrode channel. Sublayers of the hippocampus were identified from histology showing location of the dye-coated probe and electrophysiological markers, including peak theta and ripple power, theta phase shifts, gamma coherence, and density of action potentials (Karlsson and Blumberg, 2004; Lubenov and Siapas, 2009; Schomburg et al., 2014; Senzai and Buzsáki, 2017). Channels from each layer were aligned and spectrograms for each stimulation condition were averaged across animals.

To determine if stimulating SOM+ neurons reset the theta phase, we aligned the down-sampled LFP from the deepest CA1 pyramidal cell layer channel to the onset of light pulses (10ms duration) delivered at 1Hz. The LFP was averaged across 100 light deliveries and the mean envelope was calculated during a 50ms wide bin starting 250ms before and after the aligned light deliveries.

### Statistics

All statistics were performed in MATLAB 2017a or GraphPad Prism. For circular data, all phases were converted to radians and were analyzed using the Circular Statistics Toolbox in MATLAB (Berens, 2009). Linear data is presented as mean ± standard error of the mean while circular data is presented as mean ± standard deviation. Threshold for significance (α) was set to 0.05 except for the identification of photo-tagged single-units, where α was set to 0.001.

Trigger precision (r-value) during trough- or peak-targeted stimulations was tested for correlations with theta power during locomotion during the baseline period using a Pearson test to determine R^2^ value and if the slope of the line of best fit was significantly non-zero. One animal that was included for determining trigger precision was excluded from this analysis due to insufficient pyramidal cell layer data. The change in distribution and mean phase of firing (i.e., mu-values / phase preference) between baseline, trough-stimulation, and peak-stimulation conditions was compared with circular Kuiper and Watson-Williams tests, respectively. The magnitude of the phase preference (r-value) was compared with a repeated measures ANOVA followed by *posthoc* Holm-Sidak tests. The negative correlation between baseline firing rate and the change in mu during peak-targeted stimulation compared to baseline was determined using a Pearson test. Theta power (5-12Hz) was compared between baseline, trough-, and peak-targeted stimulations using a repeated measures ANOVA. Mean envelope of the aligned and averaged pyramidal cell layer LFP signal before and after a 10ms blue light pulse was compared with a paired t-test to determine if SOM+ neuron stimulation induced a theta phase reset.

## ACKNOWLEDGEMENTS

We would like to wholeheartedly thank Denise Cai for thoughtful comments and feedback on the design and implementation of this project and manuscript as well as the entire Cai and Shuman lab group for providing their support throughout the course of this project. In particular, we would like to thank Alia Abdelhameed and Helen Liu for assistance with animal training and tissue processing, and Lauren Vetere for assistance with acute silicon probe recording setup and virtual environment building. Additional thanks to Greg Hilleren for helpful discussions on circular statistics. This work was supported by a CURE Taking Flight Award (TS), and NIH grants R01 NS116357 (TS), RF1 AG072497 (TS), and F32 NS116416 (ZCW).

## AUTHOR INFORMATION

ZCW, PAP, and TS conceptualized and designed the project. ZCW performed data acquisition, and wrote the original draft of the manuscript; PAP implemented PhaSER in LabVIEW; ZCW and PAP performed formal analyses; ZCW, PAP, SIL, CK, EK, YF, and CH contributed to various aspects of the methodology including stereotaxic surgeries, tissue processing, animal training, fluorescent imaging, and custom MATLAB script writing. ZCW, PAP, SIL, CK, EK, YF, and TS iteratively reviewed and edited this manuscript. TS supervised. Corin Humphrey passed away October 2020 after making substantive methodological contributions to this project. She agreed to be an author on this project before her death.

## REFERENCES

Amilhon B, Huh C, Manseau F, Ducharme G, Nichol H, Adamantidis A, Williams S (2015) Parvalbumin Interneurons of Hippocampus Tune Population Activity at Theta Frequency. Neuron 86:1277–1289.

Aronov D, Tank DW (2014) Engagement of Neural Circuits Underlying 2D Spatial Navigation in a Rodent Virtual Reality System. Neuron 84:442–456.

Bender AC, Luikart BW, Lenck-Santini P-P (2016) Cognitive Deficits Associated with Nav1.1 Alterations: Involvement of Neuronal Firing Dynamics and Oscillations. Plos One 11:e0151538.

Berens P (2009) CircStat: A MATLAB Toolbox for Circular Statistics. Journal of Statistical Software.

Bezaire MJ, Raikov I, Burk K, Vyas D, Soltesz I (2016) Interneuronal mechanisms of hippocampal theta oscillations in a full-scale model of the rodent CA1 circuit. eLife 5.

Bures J, Fenton AA (2000) Neurophysiology of Spatial Cognition. Physiology 15:233–240.

Buzsáki G (2002) Theta oscillations in the hippocampus. Neuron 33:325–340.

Cai DJ et al. (2016) A shared neural ensemble links distinct contextual memories encoded close in time. Nature 534:115.

Chen LL, Madhavan R, Rapoport BI, Anderson WS (2013) Real-Time Brain Oscillation Detection and Phase-Locked Stimulation Using Autoregressive Spectral Estimation and Time-Series Forward Prediction. Ieee T Bio-med Eng 60:753–762.

Christenson Wick Z, Tetzlaff MR, Krook-Magnuson E (2019) Novel long-range inhibitory nNOS-expressing hippocampal cells. Elife 8:e46816.

Colgin LL (2011) Oscillations and hippocampal-prefrontal synchrony. Current opinion in neurobiology 21:467–474.

Cutsuridis V, Hasselmo M (2012) GABAergic contributions to gating, timing, and phase precession of hippocampal neuronal activity during theta oscillations. Hippocampus 22:1597–1621.

Cutsuridis V, Poirazi P (2015) A computational study on how theta modulated inhibition can account for the long temporal windows in the entorhinal-hippocampal loop. Neurobiol Learn Mem 120:69–83.

Desideri D, Zrenner C, Gordon PC, Ziemann U, Belardinelli P (2018) Nil effects of μ-rhythm phase-dependent burst-rTMS on cortical excitability in humans: A resting-state EEG and TMS-EEG study. Plos One 13:e0208747.

Donoghue T, Dominguez J, Voytek B (2020) Electrophysiological Frequency Band Ratio Measures Conflate Periodic and Aperiodic Neural Activity. Eneuro:ENEURO.0192-20.2020.

Dudok B, Klein PM, Hwaun E, Lee BR, Yao Z, Fong O, Bowler JC, Terada S, Sparks FT, Szabo GG, Farrell JS, Berg J, Daigle TL, Tasic B, Dimidschstein J, Fishell G, Losonczy A, Zeng H, Soltesz I (2021) Alternating sources of perisomatic inhibition during behavior. Neuron.

Földi T, Lőrincz ML, Berényi A (2021) Temporally Targeted Interactions With Pathologic Oscillations as Therapeutical Targets in Epilepsy and Beyond. Front Neural Circuit 15:784085.

Franklin KBJ, Paxinos G (2007) The mouse brain: in stereotaxic coordinates. Academic Press, Elsevier.

Gordon PC, Belardinelli P, Stenroos M, Ziemann U, Zrenner C (2022) Prefrontal theta phase-dependent rTMS-induced plasticity of cortical and behavioral responses in human cortex. Brain Stimul 15:391–402.

Hippenmeyer S, Vrieseling E, Sigrist M, Portmann T, Laengle C, Ladle DR, Arber S (2005) A developmental switch in the response of DRG neurons to ETS transcription factor signaling. PLoS biology 3:e159.

Holtzheimer PE, Mayberg HS (2011) Deep Brain Stimulation for Psychiatric Disorders. Annu Rev Neurosci 34:289–307.

Hyman JM, Wyble BP, Goyal V, Rossi CA, Hasselmo ME (2003) Stimulation in Hippocampal Region CA1 in Behaving Rats Yields Long-Term Potentiation when Delivered to the Peak of Theta and Long-Term Depression when Delivered to the Trough. J Neurosci 23:11725–11731.

Kanta V, Pare D, Headley DB (2019) Closed-loop control of gamma oscillations in the amygdala demonstrates their role in spatial memory consolidation. Nat Commun 10:3970.

Karlsson KÆ, Blumberg MS (2004) Temperature-Induced Reciprocal Activation of Hippocampal Field Activity. J Neurophysiol 91:583–588.

Klausberger T, Magill PJ, Márton LFF, Roberts JD, Cobden PM, Buzsáki G, Somogyi P (2003) Brain-state- and cell-type-specific firing of hippocampal interneurons in vivo. Nature 421:844–848.

Klausberger T, Somogyi P (2008) Neuronal Diversity and Temporal Dynamics: The Unity of Hippocampal Circuit Operations. Science 321:53–57.

Krause MR, Vieira PG, Thivierge J-P, Pack CC (2022) Brain stimulation competes with ongoing oscillations for control of spike timing in the primate brain. Plos Biol 20:e3001650.

Kwag J, Paulsen O (2009) Bidirectional control of spike timing by GABA(A) receptor-mediated inhibition during theta oscillation in CA1 pyramidal neurons. Neuroreport 20:1209–1213.

Laurent F, Brotons-Mas JR, Cid E, Lopez-Pigozzi D, Valero M, Gal B, Prida LM de la (2015) Proximodistal structure of theta coordination in the dorsal hippocampus of epileptic rats. The Journal of neuroscience : the official journal of the Society for Neuroscience 35:4760–4775.

Lazaro MT et al. (2019) Reduced Prefrontal Synaptic Connectivity and Disturbed Oscillatory Population Dynamics in the CNTNAP2 Model of Autism. Cell Reports 27:2567-2578.e6.

Lenck-Santini P-P, Holmes GL (2008) Altered Phase Precession and Compression of Temporal Sequences by Place Cells in Epileptic Rats. J Neurosci 28:5053–5062.

Lopez-Pigozzi D, Laurent F, Brotons-Mas J, Valderrama M, Valero M, Fernandez-Lamo I, Cid E, Gomez-Dominguez D, Gal B, Prida ML de la (2016) Altered Oscillatory Dynamics of CA1 Parvalbumin Basket Cells during Theta-Gamma Rhythmopathies of Temporal Lobe Epilepsy. eNeuro 3.

Lubenov EV, Siapas AG (2009) Hippocampal theta oscillations are travelling waves. Nature 459:534–539.

Mitra P, Bokil H (2008) Observed Brain Dynamics. Oxford University Press.

Mitra P, Bokil H, Maniar H, Loader C, Mehta S, Hill D, Mitra S, Andrews P, Baptista R, Gopinath S, Nalatore H, Kaur S (2018) Chronux.

Rahsepar B, Noueihed J, Norman JF, Lahner B, Quick MH, Ghaemi K, Pandya A, Fernandez FR, Ramirez S, White JA (2022) Theta phase specific modulation of hippocampal memory neurons. Biorxiv:2022.10.27.513992.

Rutishauser U, Ross IB, Mamelak AN, Schuman EM (2010) Human memory strength is predicted by theta-frequency phase-locking of single neurons. Nature 464:903–907.

Schatza MJ, Blackwood EB, Nagrale SS, Widge AS (2022) Toolkit for Oscillatory Real-time Tracking and Estimation (TORTE). J Neurosci Meth 366:109409.

Schomburg EW, Fernández-Ruiz A, Mizuseki K, Berényi A, Anastassiou CA, Koch C, Buzsáki G (2014) Theta Phase Segregation of Input-Specific Gamma Patterns in Entorhinal-Hippocampal Networks. Neuron 84:470–485.

Senzai Y, Buzsáki G (2017) Physiological Properties and Behavioral Correlates of Hippocampal Granule Cells and Mossy Cells. Neuron 93:691-704.e5.

Shobe JL, Claar LD, Parhami S, Bakhurin KI, Masmanidis SC (2015) Brain activity mapping at multiple scales with silicon microprobes containing 1,024 electrodes. J Neurophysiol 114:2043–2052.

Shuman T et al. (2020) Breakdown of spatial coding and interneuron synchronization in epileptic mice. Nat Neurosci 23:229–238.

Shuman T, Amendolara B, Golshani P (2017) Theta Rhythmopathy as a Cause of Cognitive Disability in TLE. Epilepsy currents 17:107–111.

Siegle JH, Wilson MA (2014) Enhancement of encoding and retrieval functions through theta phase-specific manipulation of hippocampus. Elife 3:e03061.

Sigurdsson T, Stark KL, Karayiorgou M, Gogos JA, Gordon JA (2010) Impaired hippocampal–prefrontal synchrony in a genetic mouse model of schizophrenia. Nature 464:763–767.

Skarpaas TL, Morrell MJ (2009) Intracranial stimulation therapy for epilepsy. Neurotherapeutics 6:238–243.

Somogyi P, Katona L, Klausberger T, Lasztóczi B, Viney TJ (2013) Temporal redistribution of inhibition over neuronal subcellular domains underlies state-dependent rhythmic change of excitability in the hippocampus. Philosophical Transactions Royal Soc Lond Ser B Biological Sci 369:20120518.

Steinmetz NA et al. (2021) Neuropixels 2.0: A miniaturized high-density probe for stable, long-term brain recordings. Science 372.

Stringer C, Pachitariu M, Steinmetz N, Reddy CB, Carandini M, Harris KD (2019) Spontaneous behaviors drive multidimensional, brainwide activity. Science 364:255.

Taniguchi H, He M, Wu P, Kim S, Paik R, Sugino K, Kvitsiani D, Kvitsani D, Fu Y, Lu J, Lin Y, Miyoshi G, Shima Y, Fishell G, Nelson SB, Huang Z (2011) A resource of Cre driver lines for genetic targeting of GABAergic neurons in cerebral cortex. Neuron 71:995–1013.

Wodeyar A, Schatza M, Widge AS, Eden UT, Kramer MA (2021) A state space modeling approach to real-time phase estimation. Elife 10:e68803.

Wyble BP, Hyman JM, Rossi CA, Hasselmo ME (2004) Analysis of theta power in hippocampal EEG during bar pressing and running behavior in rats during distinct behavioral contexts. Hippocampus 14:662–674.

Zelmann R, Paulk AC, Basu I, Sarma A, Yousefi A, Crocker B, Eskandar E, Williams Z, Cosgrove GR, Weisholtz DS, Dougherty DD, Truccolo W, Widge AS, Cash SS (2020) CLoSES: A platform for closed-loop intracranial stimulation in humans. Neuroimage 223:117314–117314.

